# ETDB-Caltech: a blockchain-based distributed public database for electron tomography

**DOI:** 10.1101/453662

**Authors:** Davi R. Ortega, Catherine M. Oikonomou, H. Jane Ding, Prudence Rees-Lee, Alexandria, Grant J. Jensen

**Affiliations:** Division of Biology and Biological Engineering, California Institute of Technology, Pasadena, California, USA; Howard Hughes Medical Institute, Pasadena, California, USA

**Author notes:** Corresponding author (GJJ). Membership of Alexandria is provided in the Acknowledgments.

## Abstract

Three-dimensional electron microscopy techniques like electron tomography provide valuable insights into cellular structures, and present significant challenges for data storage and dissemination. Here we explored a novel method to publicly release more than 11,000 such datasets, more than 30 TB in total, collected by our group. Our method, based on a peer-to-peer file sharing network built around a blockchain ledger, offers a distributed solution to data storage. In addition, we offer a user-friendly browser-based interface, https://etdb.caltech.edu, for anyone interested to explore and download our data. We discuss the relative advantages and disadvantages of this system and provide tools for other groups to mine our data and/or use the same approach to share their own imaging datasets.

## Introduction

Three-dimensional electron microscopy (3D EM) techniques produce large and information-rich datasets about biological samples. In electron tomography (ET), samples are imaged as they are tilted incrementally – typically 1-2 degrees between images. The resulting tilt-series of 2D projection images can then be computationally combined into a 3D reconstruction, or tomogram, of the sample with nanometer-scale resolution. ET has both biological [1] and materials science applications [2]. ET is frequently performed on frozen samples (cryo-ET) such as intact, small cells. Cryo-ET has revealed many details about cell ultrastructures that are inaccessible by other techniques, either because they cannot be purified intact or because they are not preserved by traditional EM sample preparations [3]. Another 3D EM technique, single particle analysis, also yields 3D information about cellular complexes [4].

Biological applications of 3D EM techniques are rapidly increasing, with an explosive rise in the number of datasets published [5] and excitement about the field (e.g. [6-8]). In addition, technological advances such as increased automation for higher-throughput data collection and movie acquisition with direct detectors are increasing the information content of datasets [9, 10], which makes management of these datasets a mounting challenge [11]. At the same time, public accessibility is of critical importance [12]. 3D EM techniques, while burgeoning, are still inaccessible to most cell biologists due to the expensive equipment (several million dollars to purchase and maintain, in a customized space) and specialized expertise required. In addition, the technology is still in a phase of active development, in both hardware and software. To facilitate software development efforts, programmers need access to large and varied test datasets.

Public dissemination outlets for 3D EM datasets address two fundamentally different missions: (1) to provide curated, validated data for peer review and education [13]; and (2) to provide large quantities of possibly problematic data to facilitate biological discovery and software development. The first mission is well served by resources such as the Electron Microscopy Data Bank (EMDB) and the Cell Image Library. The EMDB, an invaluable community tool for deposition of 3D EM data [14], is part of the EMDataBank [15], a global resource for 3D EM managed by the worldwide Protein Data Bank (PDB) consortium [16]. Like its counterpart, the PDB [17], it is the standard repository for published structures, such as single particle reconstructions and subtomogram averages [18]. To encourage public access, the EMDB developed web-based visualization tools to interact with data [19, 20]. The Cell Image Library^TM^ is an open-source catalog of curated images, animations and videos aimed at disseminating cell biology to the broader public [21]. Entries include light and electron microscopy imaging, as well as correlated datasets. The resource includes datasets previously available as the Cell Centered Database (CCDB), an online repository of high-resolution, often 3D, light and electron microscopy data, including many electron tomograms [22-24].

The second mission is currently served in a more piecemeal fashion, largely by initiatives from single labs and imaging centers to release a subset of their raw datasets for public access. Unfortunately, these resources often suffer from a lack of permanence due to lapsed maintenance of published websites. Recognizing the need for a centralized public repository of the raw EM datasets from which EMDB structures are derived, in 2016 the European PDB announced a sister site to the EMDB: the Electron Microscopy Public Image Archive, or EMPIAR [25]. EMPIAR collects tilt-series related to reconstructions deposited in the EMDB. It therefore offers an ideal resource for benchmarking software with verified, published datasets, but it is not designed for large-scale releases of unpublished, problematic and/or complicated datasets: datasets must be associated with an EMDB deposition; only tilt-series can be deposited (the resulting reconstructions are available in the EMDB, but associated files such as correlated light microscopy images or digital segmentations cannot be included); and much of the metadata is entered manually [26], a daunting task for a large batch of data.

While releasing data of unverified quality may seem to be of dubious value, we would argue that it is necessary for the progress of the field. As pointed out by the developers of the CCDB, ET datasets that currently yield poor-quality reconstructions offer opportunities for developing better reconstruction methods [24]. Also, biological insights often come from unexpected places; as a single anecdotal example, years ago our lab collected electron tomograms of bacteria to study chromosome segregation and observed novel tubes inside cells; we shared the images and a cell biologist made a connection to a secretion system he was studying, allowing us together to figure out its mechanism [27].

Since 2003, our lab has collected more than 30,000 ET datasets. Each dataset consists of a tilt-series of 2D TEM projection images and the resulting 3D tomographic reconstruction, as well as additional image, video, and segmentation files. Each dataset is 1-5 GB, and the full collection adds up to ~110 TB of data. To store and curate this volume of data for internal use by our group, we developed the Caltech Tomography Database, a central repository linked to a browser-based interface for lab members to browse, search, and download data [28]. To further streamline data handling, we integrated the internal Caltech Tomography Database with an automatic processing pipeline that uploads and processes datasets as they are acquired by the microscope [28] The majority of our ET datasets come from cryo-preserved cells. They represent more than 100 unique species of bacteria, archaea, and eukaryotes and have led to dozens of publications about diverse aspects of cell ultrastructure. The nature of whole-cell imaging, though, means that these datasets are far from exhausted. While we collected them for a specific study, they contain information about many other aspects of cell biology that may be useful to other researchers.

While we have been sharing our data by publishing papers and depositing representative tomograms in the EMDB, we have also received many requests–from software developers, biologists, and EMPIAR–to share more of our data. We filled these individual requests, but wanted to explore a broader solution to enable our lab and others to share large amounts of data of unverified quality in a persistent and decentralized fashion. The approach we describe here uses a distributed peer-to-peer file network tracked by an ownerless ledger (blockchain) system. We describe how we used this method to release more than 11,000 electron tomography datasets (excluding those that are still part of ongoing studies), representing 85 species and encompassing more than 30 TB. We discuss the advantages and drawbacks of our approach, and how it can be adopted by other groups that wish to share their own datasets.

## Results & Discussion

### Approach

In recent years, decentralized cryptographic ledgers, or blockchains, have been explored as a method to securely record data (typically cryptocurrency transactions, for which they were first conceived [29]). Rather than relying on a trusted central authority, blockchains employ a security model that builds consensus from a system of distributed users, none of whom necessarily need to trust one another. Originally developed to solve the problem of double-spending, blockchain technology has since been adapted to other uses. For instance, the Republic of Georgia uses the bitcoin blockchain to record land transfer titles, one of several countries using the cryptographic ledger to improve the security of property rights [30]. In the United States, blockchains have been proposed as a way for patients to control access to their digital medical records [31, 32]. Blockchains are used by Nasdaq in the U.S. and stock exchanges in other countries to record private securities transactions [33].

In 2013, an anonymous developer announced a fork from a cryptocurrency called Litecoin to create a new cryptocurrency, FlorinCoin (FLO), whose ledger features a descriptive transaction comment line similar to that found on a traditional check. The text entered in this transaction comment is stored in the FLO blockchain along with the details of the transaction. Each comment can contain up to 528 characters [34]. In 2014, a company called Alexandria proposed to use this feature as a public record of information and developed an open source protocol termed the Open Index Protocol (OIP) [35]. They first used this protocol to record public social media status in the FLO blockchain and later, using a peer-to-peer distributed file-sharing network, they expanded the specifications of the protocol to register the metadata of videos and music in the FLO blockchain while storing the files in the peer-to-peer file-sharing network BitTorrent, allowing artists to prove ownership of these digital assets. From September 2017 to May 2018 FlorinCoin passed through a series of upgrades. It was renamed FLO, its code was updated to version 0.15 of Bitcoin (still retaining the sCrypt algorithm for proof-of-work), and the comment field was expanded to 1,040 characters. The current OIP specification (0.42) is optimized for the new FLO comment field size, encompasses a variety of data types, and uses a peer-to-peer file system called the InterPlanetary File System (IPFS) [36] to store files. File metadata is thereby cryptographically secured, and completely searchable, allowing anyone to discover and download the files from the IPFS.

We were curious to see if this blockchain-based data distribution model would be effective to openly and securely share our scientific imaging data. In the scheme, each dataset would be distributed to IPFS and its metadata recorded in the FLO blockchain. Any interested party, typically through a user-friendly front-end in their web browser, could query the blockchain for datasets of interest and retrieve them from IPFS. We called the resulting distributed database the public Electron Tomography Database - Caltech (ETDB-Caltech), and its information flow is schematized in Figure 1.

**Figure 1.**
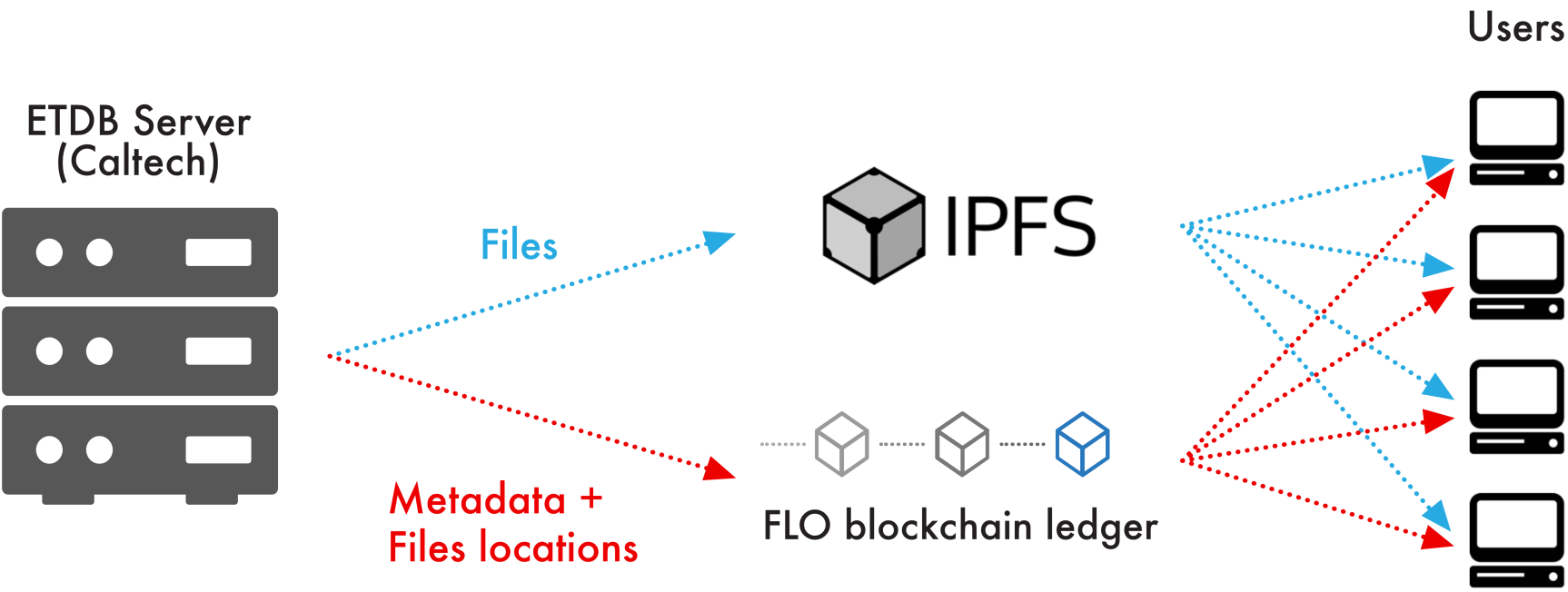
Information flow in the ETDB-Caltech file-sharing network. Datasets hosted from a local server are distributed to IPFS, a network of seeding nodes that includes the local server. The associated metadata and locations of the files are recorded in the FLO blockchain using the OIP specification. Users can query this ledger to locate and retrieve desired files from the IPFS.

We worked with Alexandria to develop a digital record type tailored to the metadata of our datasets that could be encoded easily in the FLO transaction comment. The result, Research-Tomogram, contains fields corresponding to the information we store about each dataset in our internal database. This information includes details about the user who collected the data, descriptions of the sample and its preparation, and data acquisition and processing parameters. Where appropriate, this information follows standard conventions for the 3D EM field [37]. We wrote a simple GoLang script to automatically read this information from the record in the internal lab database and translate it into an OIP Research-Tomogram record. If other groups want to adopt this approach, they can use a subset of these fields and/or add their own as necessary to match their local recordkeeping. Table 1 lists the currently available fields in the Research-Tomogram record.

**Table 1.**
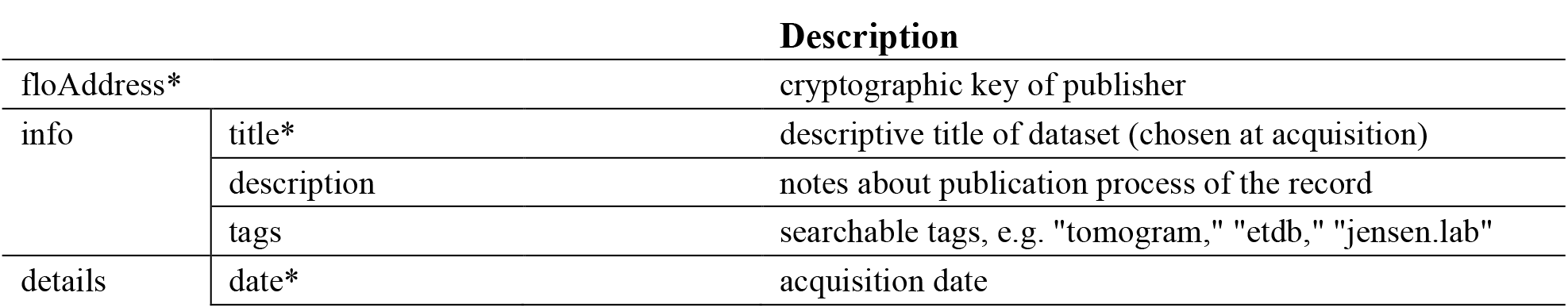
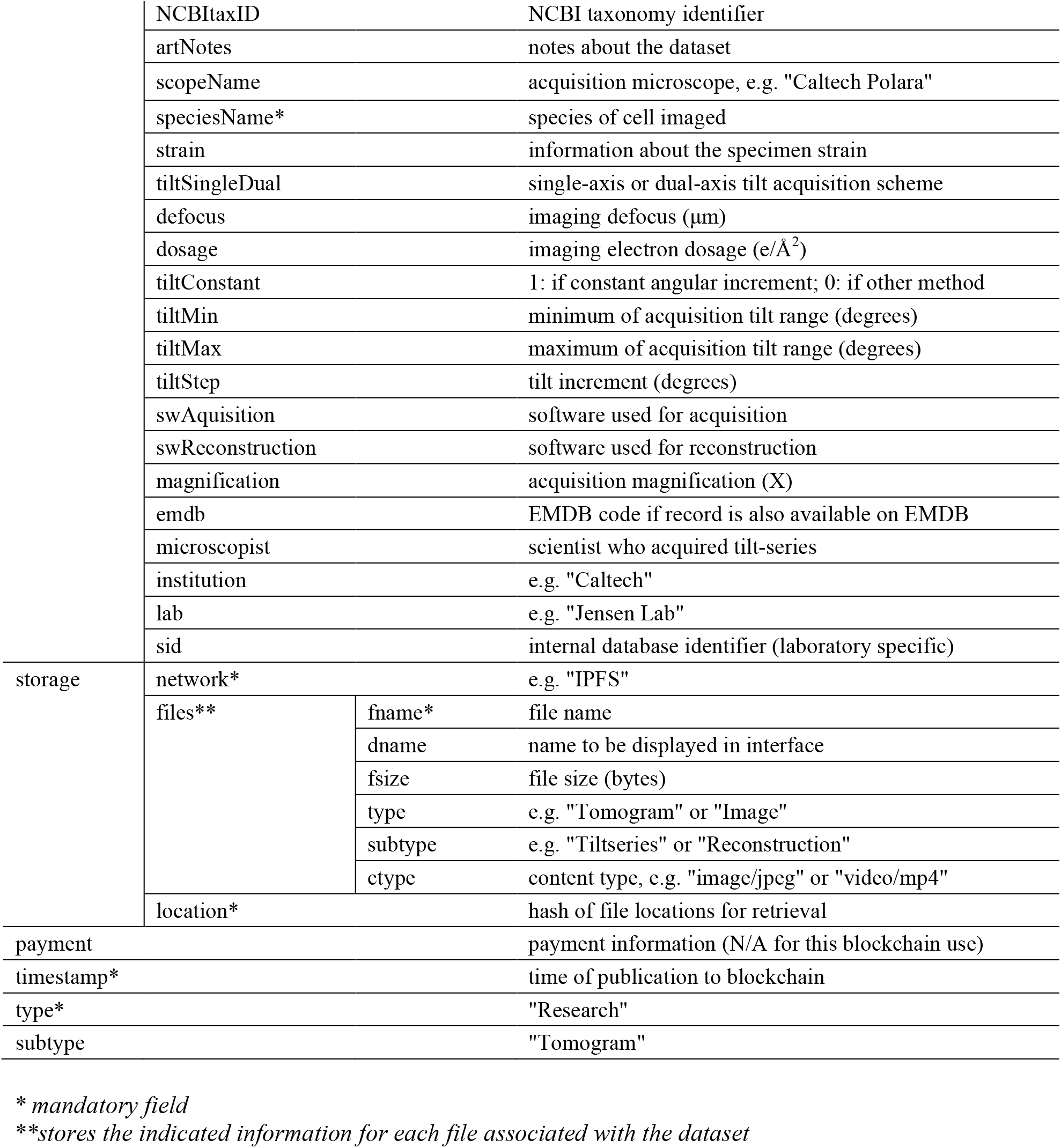
Fields in the Research-Tomogram record.

As in other peer-to-peer networks, files can be chunked and hosted from multiple nodes in the network. Users who download a file and participate in IPFS can choose to host it in this fashion for other users. This feature makes the distribution model scalable; if many users are downloading a file, multiple seeds speed up those downloads, avoiding a bottleneck from a single server. In our case, we expect relatively light file traffic, so at the current time, files are downloaded solely from our server, as in a traditional distribution model. In the rare event that a dataset is published in error, OIP offers the option of deactivating a published record. This action will not erase the metadata published in the blockchain, but the record will no longer be available to anyone using the OIP API to search the blockchain. In that case, if a user were interested in an unavailable tomogram, they would have to search the raw data in the blockchain, and hope that the files were still in the IPFS network.

There are two ways that users can download our datasets. The first is through a direct query of the blockchain and IPFS. We built a command-line application that facilitates this approach; see *Materials & Methods* for details. To increase public accessibility, we added a second route: a browser-based front-end. This graphical interface, which can be found at https://etdb.caltech.edu, provides an intuitive, interactive experience for anyone to browse ETDB-Caltech datasets, view images and videos they contain, and download part or all of each dataset. A sample dataset display page is shown in Figure 2.

**Figure 2.**
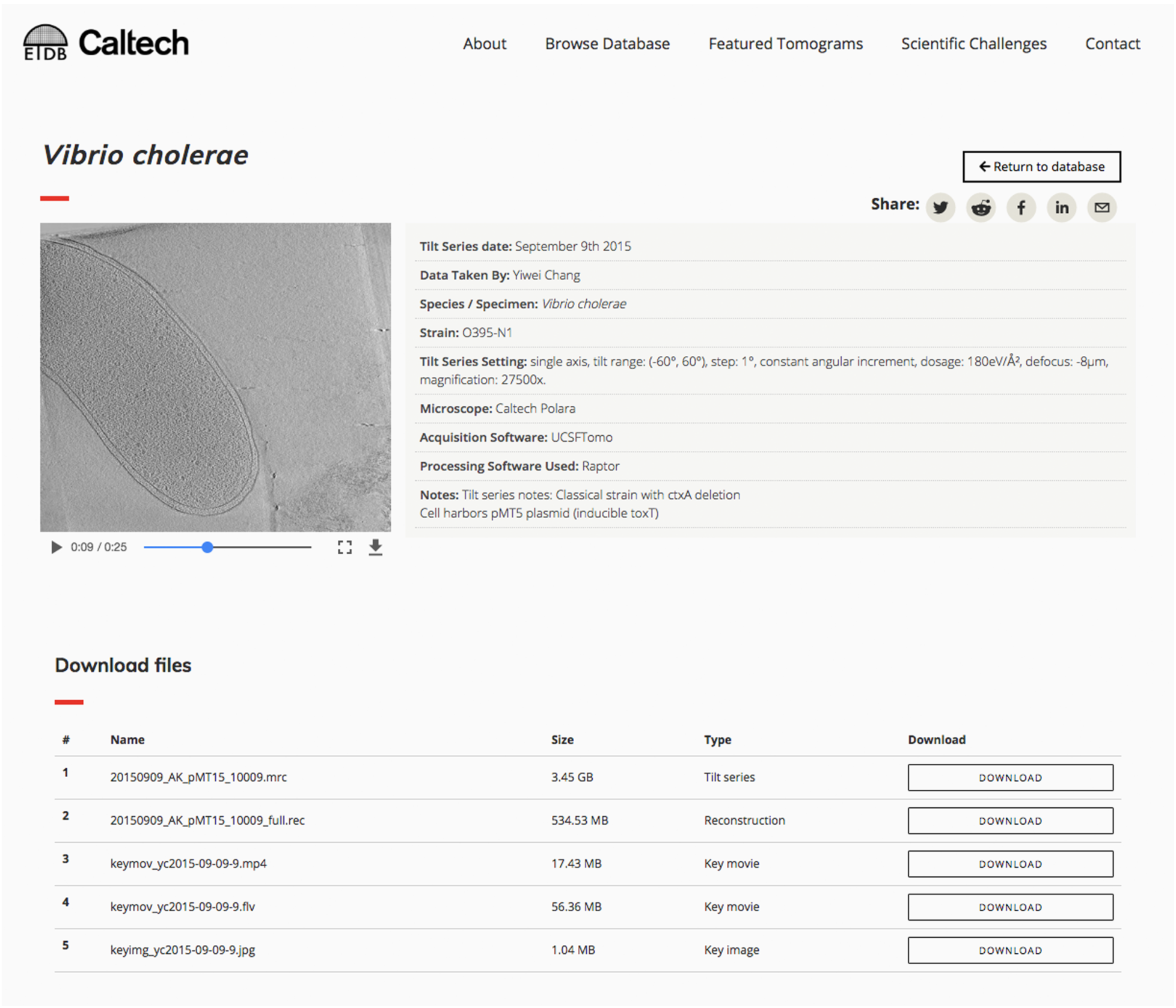
Sample entry page in the browser-based ETDB-Caltech interface. A sample electron cryotomography dataset from a *Vibrio cholerae* cell is shown. An embedded video of the reconstruction appears at left and plays automatically. The metadata is shown at right. Files associated with the dataset are listed at the bottom of the page, where they can be downloaded individually.

The ETDB-Caltech front-end offered us a chance to highlight scientific challenges for target user groups – cell biologists and software developers. We hope cell biologists will find novel features in the imaged cells, and identify those that remain mysterious. Electron tomograms contain a wealth of information, not all of which is currently interpretable; recently, for instance, we published a paper describing some of the cellular features we have observed in our electron tomograms but could not identify [38]. We hope software developers will use the released datasets to improve image-processing algorithms. In particular, we hope the availability of these datasets contributes to the development of software that can: (1) more reliably find and track the fiducial markers used for alignment in tomographic reconstruction; (2) automatically and accurately segment the boundaries of cells; and (3) automatically segment large macromolecular complexes in cells. In addition to their usefulness to experts in the field, the datasets in ETDB-Caltech may be of interest to students and the general public. To welcome these users, we designed the front-end of ETDB-Caltech to be accessible and educational, with information about the data and technology, as well as a Featured Tomograms page highlighting various features of bacterial and archaeal cells that are visible in electron tomograms (Figure 3).

**Figure 3.**
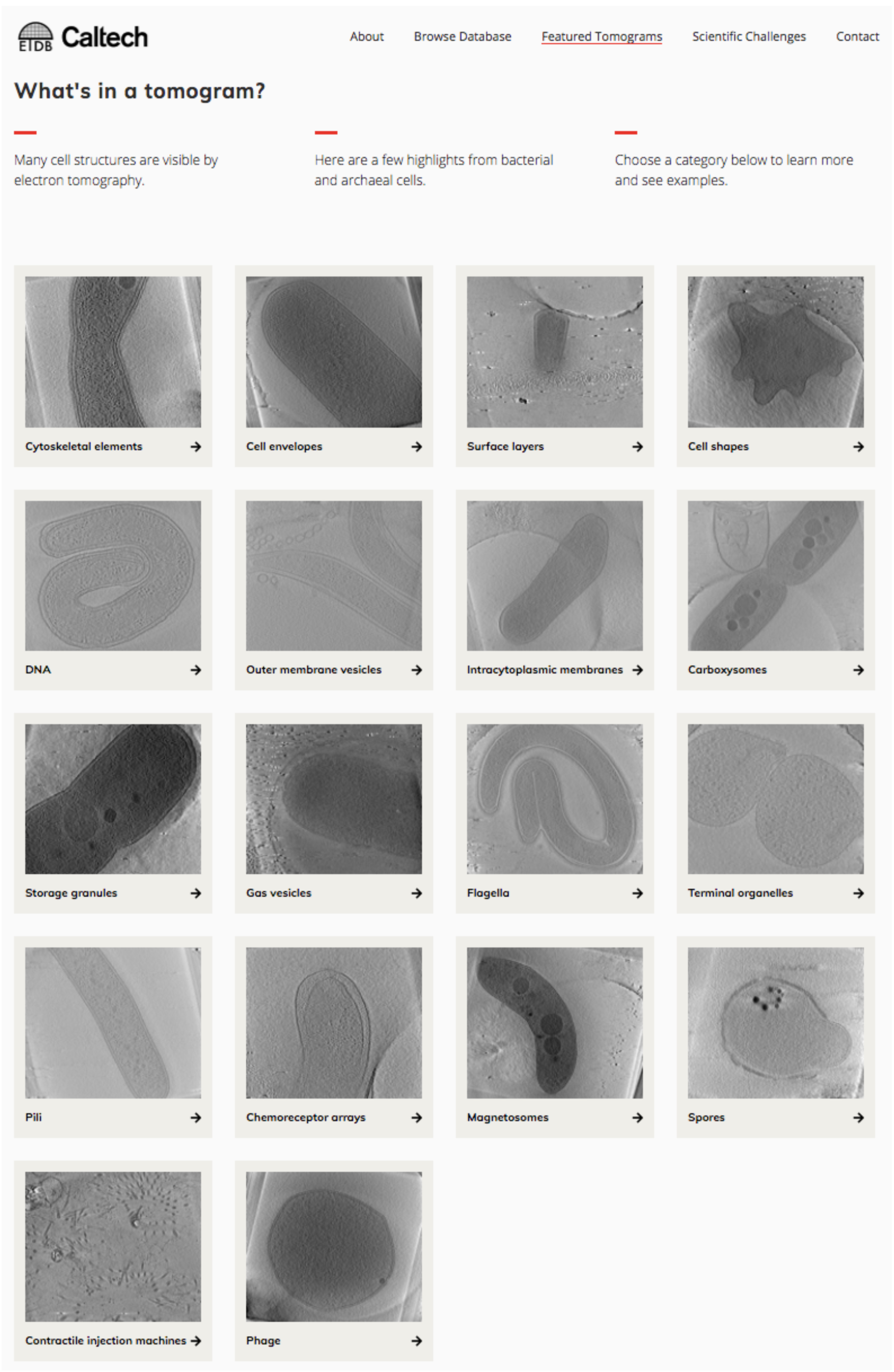
Featured Tomograms page of the ETDB-Caltech interface. Targeting students and others unfamiliar with ET data, the page highlights cellular features of bacteria and archaea visible by cryo-ET. Selecting a category takes the user to a page with a brief description of the structure and a few datasets containing examples.

### Outlook

Here we tested a new approach to publicly share a large amount of ET data. If our goal was simply to continue honoring requests from the community to make our datasets public, it would have been cheaper and easier to simply host the data from a local MySQL database, as we do for our internal group users. However, we also wanted to make a broader resource that could encompass data from many ET labs into a flexible repository that does not rely on a central authority. If ETDB is ultimately successful in enabling large-scale community data sharing, we believe it will complement (but never replace) the mission of curated repositories like EMDB and EMPIAR by providing varied datasets with a wide range of quality and content for biological and technological projects.

Compared to more centralized models of data storage, this dissemination model offers several attractive points. The first is flexibility. Multiple file types can be combined in a single OIP record, allowing, for example, light micrographs from correlative light and electron microscopy experiments and annotated segmentations to be included in EM datasets; this has been cited as a key feature lacking in some current repositories [12, 39]. Other file types from different imaging modalities can be accommodated with similar ease. The OIP specification of the Research-Tomogram record type requires few mandatory fields (Table 1). These fields can be adapted to the metadata collected by other groups, who may be using different internal databases (e.g. [40, 41]). The flip side of this flexibility is that, compared to repositories of validated datasets like EMDB/EMPIAR [26], ETDB entries may be missing information like pixel size or contain errors in metadata. This caveat should be kept in mind when using the data in further studies; information critical to interpretation should be verified with the depositor.

Another appealing feature of distributed file sharing is the distribution of storage and cost. 3D EM datasets are large, as reflected by EMPIAR, which has grown to accommodate >80 TB of stored data in 5 years [42]. These datasets are associated with only 168 studies [43]. The popularity of 3D EM methods, particularly cryo-ET [8], is growing rapidly: the number of entries in the EMDB has more than doubled over the last three years [5, 44]. There are currently more than 6,500 entries in the EMDB [44]; if each of these was associated with a similarly-sized dataset in EMPIAR, more than 3 PB of centralized storage space would be required. In a distributed distribution model, each contributing lab is responsible for storing their own data, which they presumably already do. In our case, we could have implemented the system using our existing server, which hosts our internal database, at no added cost. For extra security, we chose to keep the server with the internal database behind a local firewall and mirror the relevant datasets on an additional server outside the firewall hosting ETDB. This second server, which is larger than necessary to accommodate additional applications and future growth, cost ~US$7,000.

In addition to the local server, files should be available from other nodes of the IPFS. This ensures data persistence in the event of, for instance, a local disk failure. Of course, how well this feature works depends on whether the system is widely adopted. In addition to users hosting IPFS nodes, institutions can also easily archive ETDB data through the IPFS. The more nodes are hosting a file in the IPFS, the higher the bandwidth for users to download it; this scalability is a major feature of peer-to-peer networks. Currently, however, the IPFS is still experimental and, like many new technologies, unstable. For that reason, we serve the files in our front-end directly from the IPFS node running on our local server, not through the full IPFS peer-to-peer network. However, IPFS is in rapid development and we expect soon to update the front-end to fetch and serve the files from the IPFS. Our command line application for bulk download, ETDB-downloads, already retrieves the files from the IPFS network.

The maintenance of the ownerless ledger used to store the ETDB metadata, the FLO blockchain, depends on a distributed network of miners and users. This feature facilitates adoption as anyone can publish tomograms to the ETDB without having to seek permission from a central authority. However, as in other cryptocurrencies, miners and users have an incentive to participate in the FLO network depending on a combination of factors including the costs of hardware and electricity, and the value of FLO in the cryptocurrency market. Although FLO has been in circulation for over 5 years, a relatively long time by cryptocurrency standards, its eventual success is difficult to predict. If FLO becomes an inviable option, it may be necessary to switch to a different ledger system in the future (Ethereum, Namecoin, and Bitcoin Cash are all capable of storing text). Note, however, that metadata already published remains accessible as long as at least one copy of the FLO blockchain exists; we host one ourselves.

For us, the project took a few months to complete and the cost for the cryptocurrency transactions we used to publish 11,293 datasets was US$17.89 (see *Materials and Methods*). Most of the development effort was invested in the user interface as well as the scripts to automatically upload datasets to the IPFS and the metadata to the FLO blockchain using OIP. If other groups wish to adopt the same approach to make their data public, they would only need to slightly modify these scripts (available on GitHub, see *Materials & Methods*) to match their internal database descriptors. Our front-end code is similarly available on GitHub so that other groups can easily adapt it to taste and use it to display: (1) their own data, (2) all ETDB datasets in the IPFS, or (3) a custom subset (e.g. data from a single species or technique). In addition, individuals interested in web applications for visualization and manipulation of tomograms can use the ETDB as a distributed database of content without needing to host any tomograms themselves. Outlets (e.g. science educators) can stream tomogram videos directly from the IPFS network.

Ultimately, we believe the relationship between the ETDB and curated central repositories like the EMDB is complementary. We will continue to support the invaluable mission of the EMDB and EMPIAR in safeguarding scientific data by submitting representative curated datasets we use in our publications. We hope that the ETDB can in turn help facilitate broader releases of large batches of electron tomography data for community use. If successful, the ETDB could even be integrated into centralized repositories by their hosting an IPFS node, enhancing accessibility of the data. The flexible features of this blockchain-based, distributed scheme of data sharing may also make it useful for other types of scientific data.

## Materials & Methods

### ETDB-Caltech Distribution

The ETDB-Caltech database is fed by a MySQL database (version 14.14 distribution 5.7.21) hosted on an Ubuntu Server (Artful Aardvark kernel version 4.3.0-37). The MySQL database contains the metadata of entries from the Caltech Tomography Database [28] that have been designated for publication. Associated files are stored in a RAID6 ext4 file system. Each night, the internal server hosting the internal Caltech Tomography Database executes a script to find datasets newly edited or marked for publication and copy them to the external ETDB-Caltech server, updating the MySQL database.

The ETDB-Caltech server runs a full node of the FLO blockchain, a node of the IPFS and a MySQL database. Upon changes in the MySQL database, a custom-built GoLang script (go-etdb, available on Github: https://github.com/theJensenLab/go-etdb) makes the new files publicly accessible through the InterPlanetary File System (IPFS, version 0.4.15-dev) [36]. The IPFS daemon calculates a unique identifier to the dataset directory called a hash which is cryptographically dependent on the contents of the directory and makes the directory available to other nodes of the IPFS. This hash is combined with the metadata of each dataset and formatted according to Open Index Protocol (OIP, version 0.42) specification to create a JSON record (see Table 1). Each record generated this way is signed with a cryptographic key unique to the Jensen lab (the private key associated with public address FTSTq8xx8yWUKJA5E3bgXLzZqqG9V6dvnr) and published to the FLO blockchain by a daemon (OPId) on the server, attaching the record to the "floData" field of one or more transactions. The cost to publish the full set of 11,293 tomograms (at then-current rates of exchange) was US$17.89.

To search for ETDB-Caltech data, any user can use the cryptographic key given above to query the blockchain and retrieve matching ETDB records. This procedure is facilitated by an OIP daemon that scans and indexes the FLO Blockchain and exposes an Application Programming Interface (API) for public use. The API is accessible by a package (oip-js) deposited on the node package manager (npm). We also developed a command-line application for Unix-related environments (ETDB-downloads, manual available on Github: https://github.com/theJensenLab/etdb-downloads/blob/master/userManual.md) designed to allow users to download all or a subset of ETDB-Caltech datasets. Unlike the ETDB-Caltech website (see below), this application launches a temporary IPFS node and fetches the files from the IPFS network.

### ETDB-Caltech Interface

The front-end was built using node.js (version 9.1), react (16.2.0), webpack (4.1.1), and Twitter Bootstrap. It uses the oip-js package (https://github.com/oipwg/oip-js) to connect to an OIPdaemon Representational State Transfer (REST) API, which scans the FLO blockchain for valid OIP records and indexes them into an internal database. Currently, oip-js queries OIPdaemon for a list of records with type "Research" and subtype "Tomogram" published by our lab (the private key associated with public address: FTSTq8xx8yWUKJA5E3bgXLzZqqG9V6dvnr). In the future, queries could also search for the cryptographic keys of different groups. Alternatively, records could be retrieved by a full-node search of the FLO blockchain (available on GitHub: https://github.com/floblockchain/flo) with OIPdaemon. Files are served for download from this interface directly from the IPFS node on the ETDB-Caltech server.

The interface was designed to be easily navigable by scientists and non-scientists, and is optimized for viewing on all common web-enabled devices. We expect that in the future, some users and other labs may wish to customize this web interface. They can either copy and modify our template (available on GitHub: https://github.com/theJensenLab/etdb-react) or develop their own. While the Caltech ETDB interface displays only entries from our lab, other users may wish to build front-ends to display data from all labs sharing data using Open Index Protocol or to display only a subset of interest, for instance only those datasets corresponding to a particular species. In that case, instead of serving the files directly from the ETDB-Caltech IPFS node, those websites would use the peer-to-peer feature of the IPFS to search for the files in multiple nodes.

## Acknowledgments

We thank members of the Jensen lab for helpful comments on the ETDB interface, as well as past and present lab members (Morgan Beeby, Ariane Briegel, Yi-Wei Chang, Songye Chen, Megan Dobro, Lu Gan, Gregory Henderson, Cristina Iancu, Andreas Kjær, Zhuo Li, Alasdair McDowall, Gavin Murphy, Martin Pilhofer, Rasika Ramdasi, Jian Shi, Poorna Subramanian, Matthew Swulius, William Tivol, Elitza Tocheva, Cora Woodward, Qing Yao, Zhiheng Yu, and Elizabeth Wright who generously allowed data they collected to be made public. We also thank other lab members whose data will be published in the future. The Alexandria team is composed of Devon Read James, Amy James, Jeremiah Buddenhagen, Sky Young, Ryan Chacon and Anthony Stewart. Thanks also to past Alexandria contributors Ryan Jordan, Ryan Taylor, and Joseph Fiscella for their work on the Open Index Protocol specification. This work was made possible through the support of the National Institutes of Health (grant R35 GM122588 to G.J.J.) and the John Templeton Foundation as part of the Boundaries of Life Initiative (grant 51250 to G.J.J.).

